# Maximum water stress is decoupled from climate, traits and growth in a xeric oak

**DOI:** 10.1101/2025.08.22.666972

**Authors:** Leander D.L. Anderegg, Robert P. Skelton, Jessica Diaz, Prahlad Papper, Piper Lovegreen, Anna T. Trugman, David D. Ackerly, Todd E. Dawson

## Abstract

Widespread drought-induced forest mortality highlights the ecological consequences of climate change, yet our ability to explain, let alone predict, the spatial patterns of forest mortality remains limited.
We conducted a range-wide survey of drought stress, growth, and allocation traits in a widespread oak species (*Quercus douglasii*, blue oak) to test the predictability of stress across populations and explore how water availability and allocation mediate spatial variation in tree growth.
Across 15 sites, we found little relationship between end-of-season water availability (predawn leaf water potentials) or maximum water stress (midday water potentials) and climate or soils. Instead, water availability (within and among sites) was predicted by access to deep water resources inferred from stem water stable isotopes.
We also found a remarkable three-way decoupling of water stress, growth, and allocation to leaf tissue, which challenged a data-parameterized mechanistic plant model.
Our results reveal that deeply rooted trees can be hydrologically decouple from above-ground climate, and that the seasonality of growth, trait development and hydraulic risk are phenologically disconnected. The complicated relationship between carbon gain and hydraulic risk in seasonal environments and limited data on critical zone hydrology are key challenges to predicting drought vulnerability.

**Plain Language Summary:** Below-ground structure complicates our understanding of plant water stress. We show that below-ground complexity and highly seasonal environments can both disconnect trees from their above-ground climate and disconnect drought mortality risk from growth and trait development.

## Introduction

Anthropogenic climate change has driven more frequent and severe droughts, with catastrophic ecological consequences such as regions-scale forest mortality (Allen *et al*. 2010; Brodribb *et al*. 2020; Hammond *et al*. 2022). Two decades of intense ecophysiological research to understand how droughts drive forests over the brink has revealed that failure of the hydraulic system, interacting with disruptions to carbon metabolism—particularly for defense against biotic agents—drives tree mortality during drought (Adams *et al*. 2017; Brodribb *et al*. 2020). This insight has sparked a focus on plant hydraulics as the key to understanding and predicting spatial patterns of forest mortality (Tai *et al*. 2016). However, our ability to hindcast which trees will die (in space and time) remains extremely limited (Anderegg *et al*. 2016; Benito Garzón *et al*. 2018; Trugman *et al*. 2021; Venturas *et al*. 2020).

Our inability to predict mortality may be driven either by a poor understanding of water supply (i.e. how meteorological conditions translate into soil moisture limitation), or by unappreciated variation in the biological factors mediating how supply limitation and atmospheric demand result in physiological stress. The first issue is the persistent challenge of translating meteorological inputs into soil moisture supply over a tree’s functional rooting depth (Anderegg *et al*. 2013). Even as our gridded climate products improve and multiply, our understanding of tree-level critical zone hydrology (from the top of unweathered bedrock to the soil surface) remains quite limited (Callahan *et al*. 2022; Dawson *et al*. 2020; Fan *et al*. 2019; Hahm *et al*. 2022; Rempe & Dietrich 2018). As a consequence, translating meteorological drought into plant available water remains challenging, particularly in geologically complex landscapes.

Water balance metrics integrating water supply and demand are the go-to climate or meteorological variables for understanding ecological drought or spatial variation in water stress (Berner *et al*. 2017; Ledo *et al*. 2018; Mitchell *et al*. 2016; Stephenson 1998, 1990). These range from simple ‘moisture deficit’ (MD, precipitation minus potential evapotranspiration) to modeled ‘climatic water deficit’ (CWD, potential evapotranspiration minus modeled actual evapotranspiration) that estimate the mismatch between water availability and demand. However, all of these approaches require substantial assumptions about soil water holding capacity. For CWD, this involves a simplified soil hydrological model, usually based on large-scale gridded soils data of 1-3m depth, and a model of actual evapotranspiration constrained by this soil bucket (Abatzoglou *et al*. 2018; Flint *et al*. 2013). These soils data typically not address deeper soil layers and are often quite coarse and inaccurate outside of agricultural areas (Terribile *et al*. 2011). Plus, our understanding of plant rooting depth remains limited (Fan *et al*. 2021; Jackson *et al*. 1996). Indeed, rock water beyond the classic ‘soil’ has proven a major water source for many forests globally (McCormick *et al*. 2021). Thus, our tools for understanding climatic or meteorological water limitation drastically simplify the complex Critical Zone hydrology that controls water supply for trees. Empirically, xylem stable water isotopes can reveal whether plants are tied to shallow soil water sources that are evaporatively enriched in deuterium (^2^H) or ^18^O or are accessing isotopically depleted water sources deeper in the Critical Zone (Ehleringer *et al*. 1993; Hahm *et al*. 2020; Sprenger *et al*. 2016). Meanwhile, predawn plant water potential (Ψ_PD_), when a tree has equilibrated with the soil, provide an empirical measurement of plant available water. Ideally, gridded water balance products would capture water availability over the functional rooting depth as inferred from time- and resource-intensive stable water isotope measurements and would predict plant water availability as measured by predawn water potentials. However, it remains an open question whether water balance metrics capture supply limitations in deep rooted species growing in geomorphically complex systems.

The second major issue for predicting drought mortality is translating supply limitation and atmospheric demand into maximum plant water stress, a process complicated by spatial variation in plant physiology. In a species with relatively fixed xylem vulnerability to embolism (e.g. *Pinus sylvestris* (Lamy *et al*. 2013) or *Quercus douglasii* (Skelton *et al*. 2019)), the minimum plant water potential (Ψ_min_, proxied by midday water potentials Ψ_MD_ during dry periods), determines a trees hydraulic risk through proximity to hydraulic damage thresholds such as xylem P50 or P88 (xylem water potential that causes 50% or 88% xylem embolism, often associated with increased risk of mortality (Adams *et al*. 2017; Hammond *et al*. 2019; Sapes & Sala 2021)). Ψ_MD_ is determined by environmental factors (soil water supply/Ψ_PD_ and atmospheric demand through vapor pressure deficit or VPD), as well as plant hydraulic efficiency that can vary substantially within species (Anderegg & HilleRisLambers 2016; Martinez-Vilalta *et al*. 2009).

In particular, allocation to evaporative leaf area versus conductive stem area (A_l_:A_s_ ratio) is a key physiological knob with which trees can balance hydraulic demand (A_l_) with hydraulic supply (A_s_) (Mencuccini *et al*. 2019b; Mencuccini & Grace 1995; Sanchez Martinez *et al*. 2020; Trugman *et al*. 2019). A_l_:A_s_ can vary within-species either through ecotypic/genetic differences among populations or plastic/acclimatory trait responses to environmental cues. In contrast to most traits, the majority of the global variation in A_l_:A_s_ is found within species (Anderegg *et al*. 2022; Sanchez Martinez *et al*. 2020). Decreasing leaf area per unit stem area helps plants avoid extremely negative Ψ_MD_ by increasing leaf area-specific hydraulic efficiency (Martinez-Vilalta *et al*. 2009; Whitehead & Jarvis 1981). While many within-species trait-climate relationships have proven esoteric (Anderegg 2023), intraspecific variation in A_l_:A_s_ in response to water limitation has emerged as a consistent pattern across studies and systems (Anderegg & HilleRisLambers 2016; Anderegg *et al*. 2021; Martinez-Vilalta *et al*. 2009; Mencuccini & Bonosi 2001; Rosas *et al*. 2019). Thus, variable allocation to leaf area may be a key acclimatory mechanism driving within-species variation in hydraulic architecture (Anderegg *et al*. 2023), with profound consequences at both the organismal and ecosystem level (Mencuccini *et al*. 2019a; Quetin *et al*. 2023). From a carbon economy perspective, biomass-based traits such as the mass of leaves relative to the mass of stems (M_l_:M_s_) are an alternative basis for quantifying leaf:stem allocation, and also show relationships with water availability (Ledo *et al*. 2018; Poorter *et al*. 2012), though they have been less studied from a water stress perspective than A_l_:A_s_.

Blue oak (*Quercus douglasii* Hook & Arn.) is an ecologically and culturally important foundation species that dominates low elevation woodlands and savannas across California, U.S.A. This deciduous, deeply rooted species is valued for dendroclimate reconstructions because of its high growth sensitivity to precipitation (Stahle *et al*. 2013), making it an ideal focal species for studying spatial variation in drought stress as water limitation appears the dominant constraint to performance. Blue oak also experienced substantial mortality during the California 2012-2016 drought (Das *et al*. 2019), though only ∼30% of the spatial variation in mortality could be explained by climate, drought severity, or groundwater depletion (Brown *et al*. 2018). Variation among populations in hydraulic damage thresholds (leaf and stem P50, or the xylem water potential that causes 50% embolism) appears limited in blue oaks (Skelton *et al*. 2019), which suggests that spatial variation in hydrology or variation in exposure to drought stress (due to trait shifts such as decreasing A_l_:A_s_ that avoid drought stress) is likely key to predicting blue oak mortality (McLaughlin *et al*. 2020).

We measured end of dry season soil moisture availability (Ψ_PD_), maximum midday water stress (Ψ_MD_), stable water isotope tracers of functional rooting depth, leaf and allocation traits, and growth rates across the geographic range of blue oak and asked: 1) can we explain the spatial variation in water availability and maximum drought stress across the landscape using climate, soil and isotopic information? 2) how do allocation traits, particularly A_l_:A_s_ vary as a function of climate or soil moisture, and do they feed back to mediate maximum water stress? 3) do individual tree growth rates correspond to site climate, observed soil moisture limitation, observed maximum water stress, or variation in traits? Finally, we used a mechanistic tree hydraulics model to integrate meteorology, our observations of supply limitation and observed traits to explore the effects on hydraulic risk and carbon gain.

We predicted supply limitation (Ψ_PD_) would be broadly correlated with climatic water balance across sites, but that climate would explain substantially less variation in maximum plant water stress (Ψ_MD_) due to changes in allocation and stomatal closure at the driest sites. We also hypothesized that observed plant water potentials would be a better predictor of both allocation traits and growth than climate or soils data, with water availability explaining allocation traits and midday water stress and the soil-leaf water potential gradient explaining growth variation. Finally, we hypothesized that a mechanistic plant hydraulics model would facilitate the integration of drought exposure and allocation traits to explain growth variation based on modeled carbon gain.

## Methods

### Study System

We sampled 15 sites across the geographic and climatic distribution of blue oak (*Quercus douglasii* Hook & Arn.) (Fig. 1a). Blue oak inhabits a wide range of precipitation and potential evapotranspiration (Fig. 1b), typified by Mediterranean-type hot, dry growing seasons (April to September) with most precipitation falling December to March.

**Figure 1:**
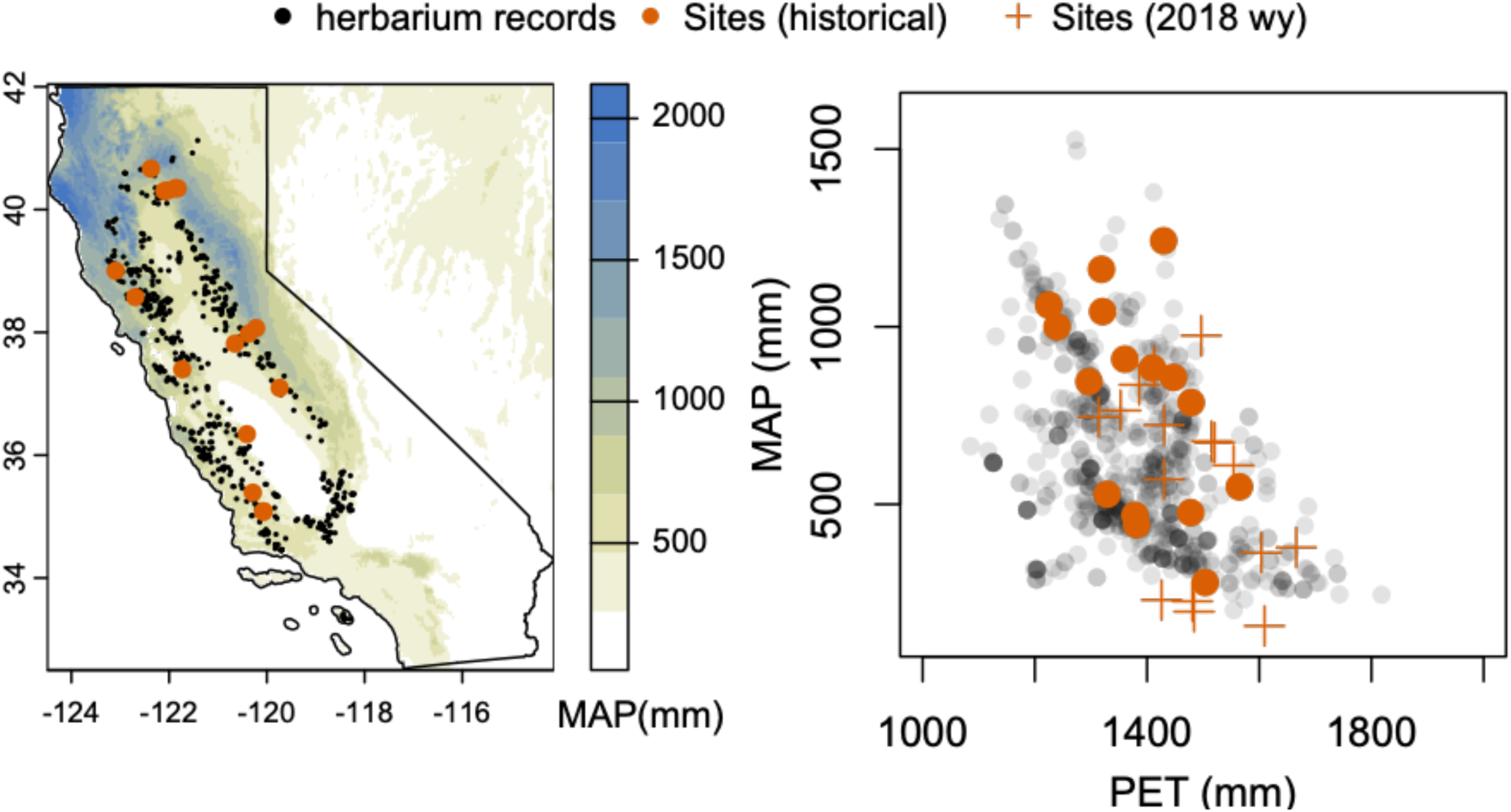
a) The geographic distribution of blue oak (Quercus douglasii) in California, USA spans a large range of mean annual precipitation (MAP in mm, from TerraClimate (Abatzoglou et al. 2018)). Black points show locations of herbarium specimens (Baldwin et al. 2017), sampled locations are shown as orange points. b) Sampled locations span the vast majority of the mean annual precipitation and potential evapotranspiration (PET) space inhabited by blue oak. The 2018 water year (crosses) when samples were collected was substantially hotter and drier than the 1981-2010 historical period (filled points).

At the end of the 2018 growing season from Sept 17^th^ to Oct 3^rd^, when trees were experiencing the annual minimum soil moisture prior to leaf senescence, we sampled predawn and midday leaf water potentials, allocation traits, leaf gas exchange, and tree cores for water stable isotope and growth analysis. We prioritized visiting all sites as quickly as possible for comparability of water potentials, at the cost of ancillary measurements at some sites. We measured water potentials at all 15 sites, but allocation traits at only 12 sites, growth and isotopes at 11 sites, and leaf gas exchange on a subset of trees at 7 sites. Very little precipitation fell (<15mm or <2% of total precip) at any site during the four months preceding our sampling. Thus, we achieved uniquely consistent antecedent drought conditions for all study sites.

### Leaf water potential

We quantified moisture availability with predawn leaf water potential (Ψ_PD_) sampled between 3am and first light (local time). Assuming limited nighttime transpiration and full equilibration with the soil, Ψ_PD_ reflects the root zone-integrated soil matric potential. We measured 3-8 leaves from each of 5-7 trees per site (mean 5.5), severing the petiole with a sharp razor blade (for short trees) or collecting a branch >50 cm long using pole pruners and immediately cutting 2-3 leaves using a razor blade. Leaves were immediately wrapped in tinfoil and a lightly damp paper towel (to limit evaporation without rehydrating the leaf) and measured in a Scholander-type pressure chamber (PMS Instruments, Corvallis, OR, USA) within 1 min of cutting. A ‘modified petiole’ (Rodriguez-Dominguez *et al*. 2022) was cut using a razor blade for leaves with short petioles.

We then measured minimum leaf water potential at midday (Ψ_MD_) between 12:30 and 2pm to estimate maximum water stress. The sampling year (2018) was drier and hotter than average (Fig. 1b), so end-of-season Ψ_MD_ was a reasonable estimate of long term Ψ_min_ values. We calculated ΔΨ (the water potential gradient driven by transpiration) as Ψ_PD_ - Ψ_MD_.

For a subset of 2-4 trees at 6 sites, we also measured midday stem xylem potential (Ψ_MD_xy_) by wrapping leaves in tinfoil, damp paper towels, and plastic bags to stop transpiration >20 mins prior to leaf collection. Ψ_MD_xy_ was on average 8% less negative than Ψ_MD_. We calculated Ψ_MD_xy_ for all trees assuming an 8% offset from Ψ_MD_ (i.e. that 8% of total resistance was between the branch xylem and leaf lamina). We used the species average P50 values for leaves (-3.88) and stems (-4.47) from Skelton et al. 2019 to calculate Hydraulic Safety Margins (HSM_leaf_ = Ψ_MD_ – P50_leaf_, HSM_stem_ = Ψ_MD_xy_ – P50_stem_). Because HSM_leaf_ and HSM_stem_ are linear transforms of Ψ_MD_ we report statistical relationships only for Ψ_MD_.

### Trait measurements

At 12 of the 15 sample sites, we collected three terminal twigs from the sun exposed mid- to upper-canopy of five trees using pole pruners. Terminal twigs included the current year (2018) stem growth, identified by previous year bud scar, and all subtending leaves, thus represent one year’s investment in branch and leaf tissue. Branches were rehydrated using the ‘partial rehydration’ method (Pérez-Harguindeguy *et al*. 2013) and stored in a cooler or refrigerator for at least 12 hours before processing. In the lab, leaves were removed (with petiole), weighed for their wet mass (M_l_wet_), and scanned on a flatbed scanner. Images were analyzed with ImageJ (Schneider *et al*. 2012) to calculate total twig leaf area (A_l_), and total leaf area was divided by leaf number for average leaf size. Stem basal diameter underneath the bark was measured at 2-5 orientations using digital calipers and used to calculate stem cross sectional area (A_s_). Total twig A_l_ was divided by stem A_s_ to calculate branch A_l_:A_s_ ratio. Stem length was also measured with calipers or a ruler. Leaves and stems were then oven dried at 60°C for 72 hours, and their dry mass measured (M_l_, M_s_). Leaf mass per area (LMA) was calculated as M_l_ / A_l_, leaf mass to stem mass ratio (M_l_:M_s_) by dividing leaf dry mass by stem dry mass, and leaf dry matter content (LDMC) by dividing leaf dry mass by leaf wet mass (M_l_/M_l_wet_). Because samples were not processed until after the water potential campaign, leaf wet mass was suspect in the oldest samples so we excluded LDMC values from samples that were stored longer than 1.5 weeks from analysis, resulting in exclusion of LDMC values from four sites.

### Stable water isotopes

Xylem water oxygen (8^18^O) and hydrogen (8^2^H or δD) stable isotope composition was measured at 11 of the study sites on 5 trees (7 trees at two sites, 3 trees at one site). Tree cores ∼4 cm long were extracted at breast height, rapidly placed in pop top scintillation vials (after removing bark and cambium), wrapped in parafilm and stored in a cooler until frozen in the lab. Water was cryogenically extracted (Ehleringer *et al*. 2000). The oxygen isotope composition of extracted water was determined through mass spectrometry by CO_2_ headspace equilibration using a Gas Bench II (GB, ThermoFinnigan) connected to a Delta Plus XL mass spectrometer (ThermoFinnigan, Bremen, Germany) at the Center for Stable Isotope Biogeochemistry (CSIB), University of California, Berkeley, CA. The hydrogen isotope composition of samples was determined via injection into a H/Device (HDEV, ThermoFinnigan, 30 Bremen, Germany) coupled to a Delta Plus mass spectrometer (ThermoFinnigan), also at CSIB. All isotopic compositions are reported in delta (δ) notation in parts per thousand (‰) relative to the V-SMOW standard

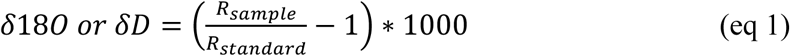

Where R = ^18^O/^16^O or D/H.

Monthly meteoric water isotope values were estimated for each site using the Online Isotopes in Precipitation Calculator, version OIPC3.2 (Bowen *et al*. 2005; Welker 2000). The evaporative deuterium enrichment of each sample relative to each site’s local meteoric water line was used to calculate ‘line conditioned’ excess (lc-excess, Landwher & Coplen 2004). Average monthly precipitation-weighted meteoric water isotopic values for the sampling year (δD_precip_, δ^18^O_precip_) were calculated for each site.

### Growth

Growth was estimated using two methods. First, we measured the branch length of terminal branches collected for trait measurements (12 sites) as a metric of 2018 branch growth. Second, we saved the tree cores collected for water isotopes after cryogenic extraction, sanded them and measured the most recent five years of ring widths using calipers and a dissecting scope to estimate recent stem growth. We translated radial growth into basal area increment (BAI) based on tree DBH. We then corrected for size-related BAI trends by calculating each tree’s growth relative to the maximum observed growth at that DBH across all sites. We fitted a quantile regression to the 90^th^ percentile BAI∼DBH relationship, and then calculated each tree’s size-standardized growth as the ‘percent of maximum BAI’ (% max BAI) or the observed BAI divided by the 90^th^ quantile BAI for the tree’s DBH.

### Climate and meteorological data

Both long term climate normals (1981-2010) and meteorological data from the 2017-2018 water year (October 1, 2017 to September 30, 2018) were extracted for each site from TerraClimate (Abatzoglou *et al*. 2018). TerraClimate has ∼4km resolution monthly climate and water balance values for the global terrestrial surface from 1958 to 2020 (updated periodically). As potential historical climate predictors (1981-2010) of plant water potential, traits, and growth, we extracted site mean annual precipitation (MAP); potential evapotranspiration (PET, based on the Penman Montieth approach); actual evapotranspiration (AET using the Thornthwaite-Mather water balance method with a single bucket model on a monthly timestep using 0.5° extractable soil water storage capacity data from (Wang-Erlandsson *et al*. 2016)); climatic water deficit (CWD = PET – AET); mean annual soil moisture (Soil_mean_ from bucket model); incident shortwave radiation (SWR); maximum temperature of the warmest month (T_max_); minimum temperature of the coolest month (T_min_); maximum annual, spring average and growing season average vapor pressure deficit (VPD_max_, VPD_spr_, VPD_gs_). We also included potential predictors from the 2017-2018 water year (termed ‘water year meteorology’ throughout), including PPT, PET, AET, CWD, T_min_, T_max_ and Soil_mean_ from the entire water year (2018 wy [variable]); plus seasonal variables including growing season PPT, VPD, and Soil_min_; dormant season PET; and spring T_max_ and T_min_. These variables were selected based on *a priori* expectations for variables that should influence growth and water stress with the inclusion of a few seasonal variables that were not highly correlated with annual variables. To avoid model overfitting and collinearity, we only fit one climate/meteorological predictor in any model.

### Soils data

We extracted plant available water (PAW), available water storage (AWC), and minimum bedrock depth for each site from the Soil Survey Geographic Database (SSURGO). The data was collected at scales ranging from 1:20,000 to 1:24,000. Both AWC (0-100 cm) and minimum bedrock depth were taken from the soil map unit composition. In two sites, where minimum bedrock depth was not available, we used average data from the dominant soil series. PAW (cm) was pulled from the dominant soil series in the soil map unit.

### Leaf gas exchange

For 1-4 trees at 7 sites, 16 total trees, we estimated stomatal conductance (g_s_) and photosynthesis (A_s_) as a function of leaf water potential at midday. Using a LI-6800 gas exchange system (LI-COR Biosciences, Lincoln, NE, USA), we collected spot measurements of g_s_ and A_s_ by setting the chamber to ambient temperature, CO_2_ concentration and relative humidity, and light to 1500 μmol m^-2^ s^-1^. Measurements were made after the conditions in the chamber stabilized (∼1min) but before stomata began to react to the chamber environment (typically 3-5 mins) and corrected for leaf area. Water potential was then measured on a leaf immediately adjacent to the measured leaf. At one site, repeated gas exchange and water potential measurements were conducted from 7 a.m. to 12:00 p.m. on one tree.

### Mechanistic model simulations

We used the HOTTER model (the **H**ydraulic **O**ptimization **T**heory for **T**ree and **E**cosystem **R**esilience model (Quetin et al 2023), a physiologically-based tree model with a realistic representation of gas exchange (B. Eller *et al*. 2018) and a detailed representation of plant hydraulics (Trugman *et al*. 2018) to quantify spatial variations in tree water status, hydraulic stress, and carbon gain across gradients in climate and plant traits in the continental United States (US). A detailed model description can be found in Quetin et al. (2023).

HOTTER model forcings include atmospheric [CO_2_], temperature, VPD, and soil water potential. In addition, the model requires inputs or allometric equations for tree leaf area, and tree size. The hydraulic and photosynthetic dynamics of the model are primarily controlled by three plant physiological traits: xylem P50, the conductivity of the roots, xylem, and petioles (K_max_), and the maximum rate of carboxylation (V_c,max_), a trait which is sensitive to temperature. Outputs include daily tree-level transpiration, gross and net carbon assimilation, canopy conductance, water potentials, and percent loss of conductivity (PLC) for roots, xylem, and leaves, assuming 12 hours of daylight for photosynthesis and 24 hours for respiration.

We used species mean traits from (Anderegg *et al*. 2023; Skelton *et al*. 2019) (Vcmax25 = 20 umol m-2 s-1; P50 = -4.3 MPa). Kmax was estimated from early growing season Knative (Kmax = 7600 mmol m-2 s-1 MPa-1). Leaf area was parameterized scaling branch A_L_:A_S_ to the tree level. Observed Ψ_PD_ was used as the soil water potential forcing, and temperature and VPD were derived from 1-hourly, 1.5-km resolution dynamically downscaled historical climate data using the Weather Research and Forecasting Model (WRF). Climate data was aggregated daily, using the 80^th^ percentile temperature and corresponding VPD as representative of the daily average conditions for the period of Sept 17-27, the same dates as the water potential sampling campaign. These dates spanned a wide range of climate conditions, as a heatwave drove an approximate doubling of VPDs between the beginning and end of the sampling period for all sites.

To estimate variability in plant performance during the sampling period, we performed separate experiments on the median, most, and least stressful day during the sampling period (as defined by VPD). We ran separate simulations for each tree with tree-specific leaf area and soil water, site-specific climate, and species-specific hydraulic and photosynthetic traits. Additionally, to examine the uncertainty in how leaf allocation influences carbon and water dynamics, we repeated these experiments but reduced tree leaf area by half.

### Analysis

We averaged all attributes to tree for analysis (87 total trees, fewer for traits, water isotopes and growth). We then sought to explain spatial variation in plant water potentials (Ψ_PD_, Ψ_MD_, ΔΨ), plant traits (A_l_:A_s_, M_l_:M_s_, leaf size, LMA, LDMC), xylem stable water isotope composition (δ^18^O, δD, and lc-excess), and growth (branch length, tree height, % max BAI) using information-theoretical based model selection, in the R statistical environment (version 4.3.1, (Team 2016)). We fit linear mixed models relating each response variables to each of the 20 climate or meteorological predictors (described above) with a random intercept for site, using the lmer() function in the ‘lme4’ package (Bates *et al*. 2015). We then used AICc (Akaike’s Information Criterion corrected for small sample sizes) to select the most parsimonious model of the 21 possible models (20 climate predictors plus a null model). Predictors were considered significant if they improved model AICc by 2 or more units over the null model. We used t-tests based on Satterthwaite’s approximate degrees of freedom to determine the p-value of significant climate predictor, using the ‘lmerTest’ R package (Kuznetsova *et al*. 2016).

We also used linear mixed effects models with a site random intercept to test whether: 1) xylem water stable isotopes explained tree-to-tree variation in plant water potentials, 2) water potential explained growth (either % max BAI or stem length), 3) whether traits predicted growth, and 4) whether HOTTER output (PLC, GPP or NPP) predicted growth. Functionally, these models tested whether, on average, there was a significant relationship among trees within a site. Thus, we also fit linear models using site-level means for all of the above relationships to test whether there were significant relationships among sites.

Data are available in the Dryad repository [insert link]. Analysis code is available in the on Github at https://github.com/leanderegg/BlueOakLandscape [*NOTE: For peer review, all relevant data can be found in the project Github repository].

## Results

### Predicting drought stress

Despite sampling across large geographic differences in precipitation and potential evapotranspiration (Fig. 1), and despite massive differences between sites in leaf water potential (-0.82 MPa to -3.78 MPa for Ψ_PD_, -2.02 MPa to -4.45 MPa for Ψ_MD_), neither Ψ_PD_ nor Ψ_MD_ could be predicted by long term site climate, 2018 water year meteorology, or soil characteristics (Fig. 2). The tree-to-tree variation within a site was substantial (mean within site range of -1.3 MPa for both Ψ_PD_ and Ψ_MD_), and was partially explained by tree size, with larger DBH trees having less negative Ψ_PD_ (Fig. S1). However, the between-site variation still constituted 71% (Ψ_PD_) and 51% (Ψ_MD_) of the total water potential variation (unbiased Ω^2^ estimate). The ΔΨ (Ψ_PD -_ Ψ_MD_, potential gradient caused by transpiration) increased with growing season VPD (linear mixed model p=0.011). The average leaf hydraulic safety margin (HSM, Ψ_MD_ – P50_leaf_) was 0 or even negative for a number of both climatically wet and dry sites, and some individual trees from both wet and dry sites had negative stem hydraulic safety margins (Ψ_MD_xy_ – P50_stem_) (Fig. 2d).

**Figure 2:**
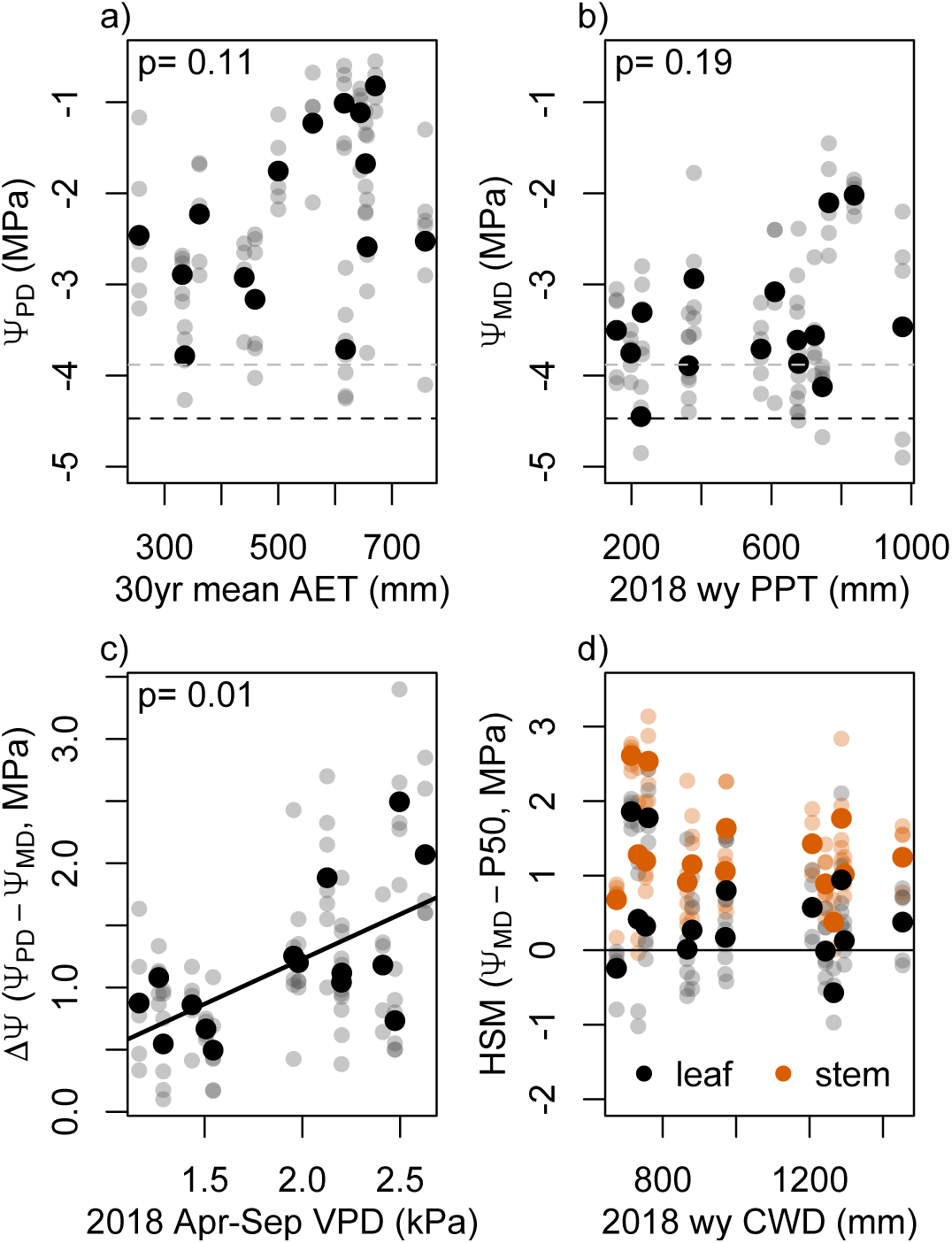
At the end of a dry growing season, neither predawn leaf water potential (a, Ψ_PD_, root zone-integrated soil moisture availability) nor midday leaf water potential (b, Ψ_MD_, maximum water stress) were predicted by site climate. (c) Transpirative potential drop (ΔΨ) increased with site growing season vapor pressure deficit (VPD). (d) hydraulic safety margin (Ψ_MD_ – P50) for leaves (black) and stems (orange). Gray points indicate individual tree average leaf water potentials, black points indicate site average leaf water potentials. Predawn and midday leaf water potentials are plotted against the best climate (1981-2010 mean) or sampling year meteorological (2017-2018 water year or ‘2018 wy’) variable identified by model selection. AET: actual evapotranspiration, PPT precipitation. p-values from linear mixed effects models with site as random intercept. Gray and black dashed lines indicate leaf and stem P50, respectively, from (Skelton et al. 2019).

Xylem stable water isotopes, on the other hand, strongly predicted leaf water potentials, both among sites and among trees within a site. Xylem water more enriched in δD and δ^18^O was associated with more negative Ψ_PD_, more negative Ψ_MD_ (smaller or negative HSMs), and a smaller ΔΨ (Fig. 3, Fig. S2). One site (Pepperwood Preserve, ‘PWD’, gray in Fig. 3) showed anomalously hydrated water potentials given its water isotopic signature, possibly due to a moderate severity wildfire one year prior to sample collection. Even with this site, leaf water potentials were significantly related to both δD and to a lesser extent δ^18^O, and without the site these relationships were highly significant (Table S2). Indeed, isotopic enrichment strongly predicted Ψ variation within site (mixed model marginal R^2^ for δD was 0.60 for Ψ_PD_, 0.35 for Ψ_MD_/HSM and 0.30 for ΔΨ) and even mores strongly between sites (R^2^ for linear model of site means was 0.84 for Ψ_PD_, 0.52 for Ψ_MD_/HSM and 0.64 for ΔΨ when excluding PWD). Site δD was strongly related to the precip-wieghted average meteoric δD_precip_ of the site, as was Ψ_PD_, suggesting that the atmospheric conditions controlling meteoric water isotope conditions were more strongly linked to end-of-season water availability than any of the climate or meteorology predictors tested (Fig. S3). However, xylem δD was still a significant predictor of Ψ_PD_ even when meteoric average δD_precip_ was included as a covariate in the linear mixed model (p<0.001 excluding PWD), and tree-level residuals from the δD_xylem_ ∼ δD_precip_ relationship were also significantly related to Ψ_PD_ (Fig. S3). The line-conditioned deuterium excess (lc-excess), which is designed to isolate the effect of evaporative enrichment on ecosystem water, was unrelated to plant water potentials.

**Figure 3:**
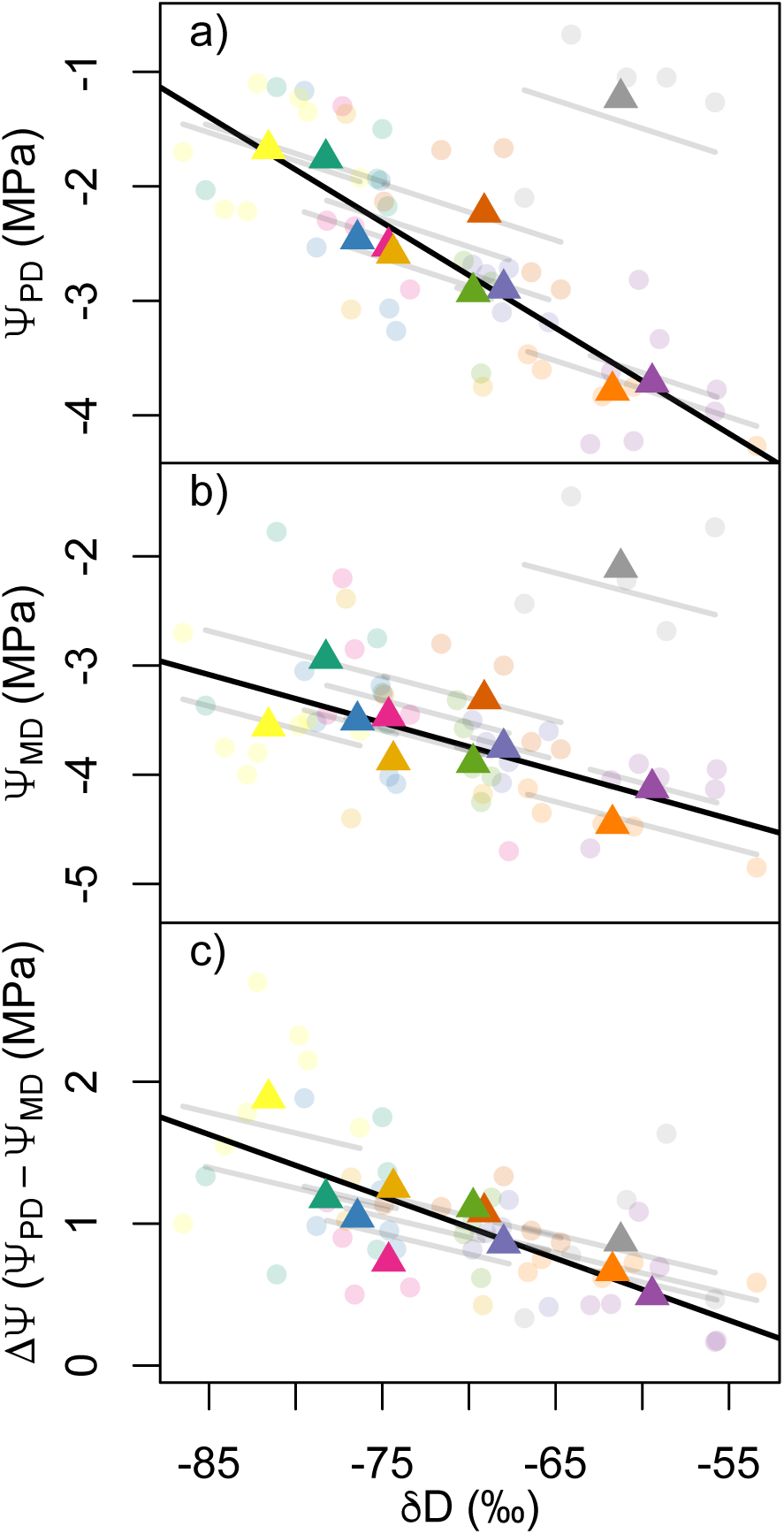
Plant water potentials were strongly related to xylem water isotopes within and among sites. Predawn leaf water potential (a), midday leaf water potential (b) and the difference between them (c) decrease with increasing xylem water enrichment of δD. Points show tree average values, triangles show site average values, colors indicate different sites, and gray lines show the trend among trees at a site. Black trend lines show statistically significant (p<0.05) fixed effects from a mixed model with site random intercepts, with the outlier burned site (gray points, triangle) excluded, and colored triangles indicate that the relationship is also significant across site means (linear model p<0.05).

Plant water potential also had profound effects on end of season leaf gas exchange. Across sites and trees within a site, midday stomatal conductance and assimilation steadily declined as Ψ decreased towards -4 MPa (Fig. 4). The point of stomatal closure coincided quite closely with the previously reported P50 of leaves of -3.88 MPa (Ψ causing 50% embolism), a threshold that was consistent across populations of blue oak (Skelton *et al*. 2019). This point of stomatal closure was also apparent in the morning time course of a single tree as its leaf water potential approached -3.88 near midday (Fig. 4). Thus, the consequences of observed water potential variation across the landscape for late season carbon gain and water loss were substantial. At two of the sampled sites, mean Ψ_PD_ was more negative than -3.7 MPa (Fig. 2), implying sustained stomatal closure and limited carbon gain at these sites for the entirety of the late growing season.

**Figure 4:**
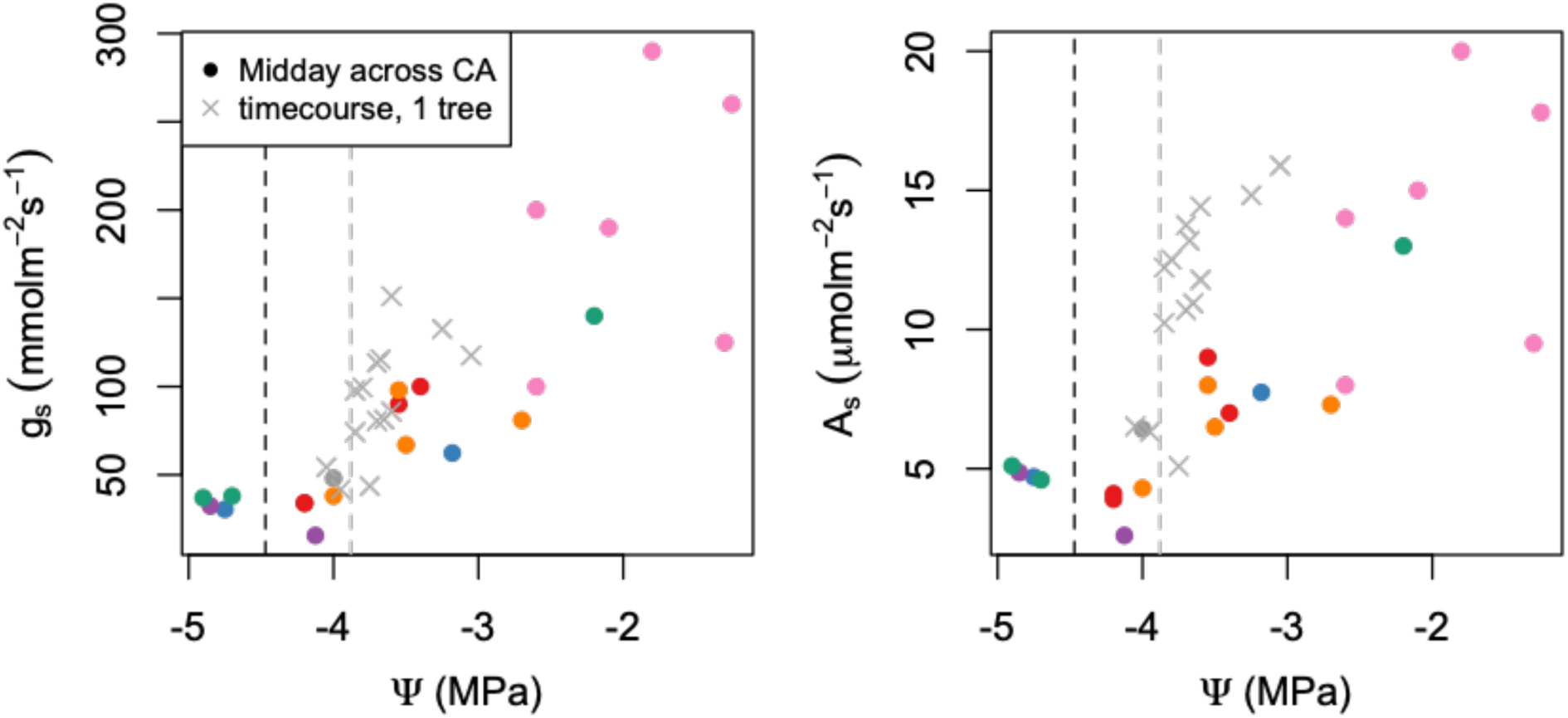
Stomatal conductance (a) and photosynthesis (b) as a function of leaf water potential. Filled circles show midday gas exchange and water potential values (color indicates different sites). Gray x’s indicate the morning time-course of a single tree as it approached midday. The gray and black vertical lines indicate the leaf and stem P50 of blue oak from Skelton et al. 2019.

### Predicting traits

All leaf and stem traits demonstrated significant site-to-site variation, ranging from 17% (M_L_:M_S_) to 58% (LDMC) of total trait variation (Ω^2^ values, Table S1). Site climate or meteorology was a significant predictor of only two traits (M_L_:M_S_ and leaf size), while soil water holding capacity predicted A_L_:A_S_ and LMA (Fig. 5). M_L_:M_S_ decreased with increasing minimum temperature of the sampling year, and leaf size increased with increasing historical precipitation. Meanwhile, A_L_:A_S_ increased and LMA decreased with soil water holding capacity (and were not predicted by any climate variables excluding soils). LDMC was not predicted by any environmental variable.

**Figure 5:**
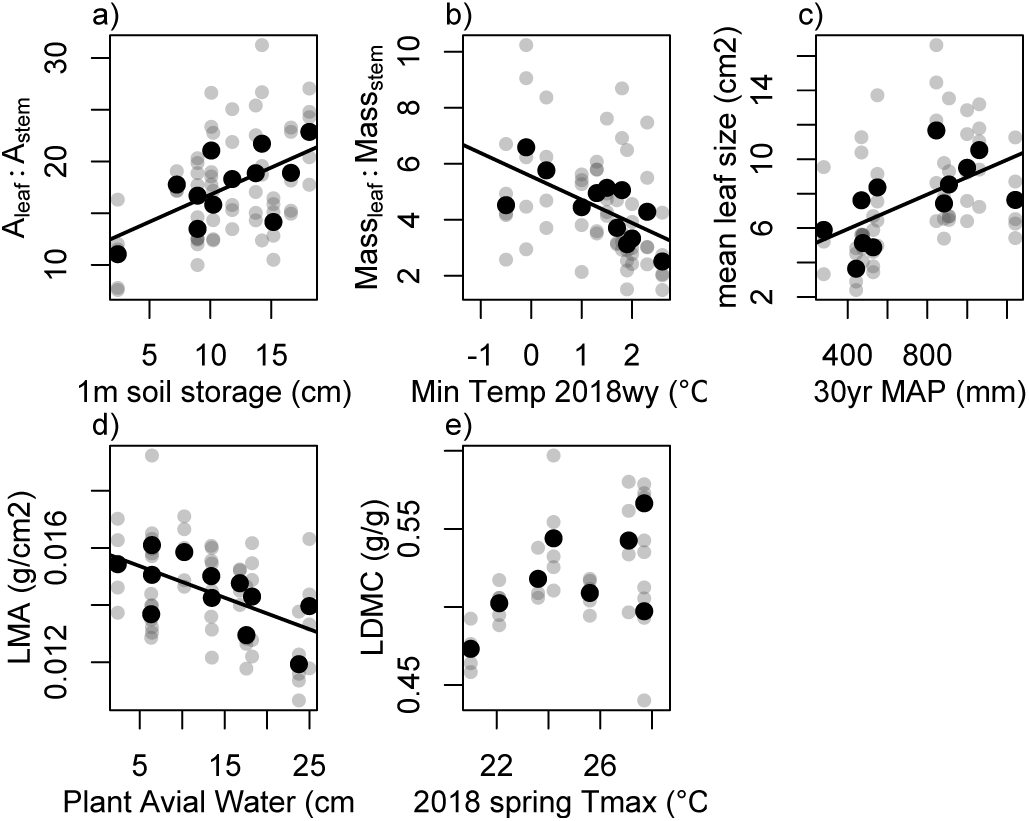
Tree-average terminal branch leaf area to stem area ratio (a, A_l_:A_s_), leaf mass to stem mass (b, M_l_:M_s_), mean leaf size (c), leaf mass per area (d, LMA) and leaf dry matter content (e, LDMC) as a function of the best climate, sample year meteorology, or soil predictor based on model selection. Black points show site means, solid trend lines show significant relationships (p<0.05).

End-of-season soil moisture and maximum water stress were almost completely unrelated to leaf and allocation traits. Ψ_PD_ was not correlated with any trait, and Ψ_MD_ was only correlated with LDMC (Fig. S4), with less negative Ψ_MD_/larger HSMs being associated with *higher* LDMC (linear mixed effects model p=0.004). Moreover, traits were largely uncorrelated with each other. A_L_:A_S_ was positively correlated with M_L_:M_S_, as well as leaf size (Fig. S4), but no other traits were significantly correlated.

### Drought and trait relationships with growth

Branch elongation and basal area growth were significantly positively correlated, but the relationship was relatively weak (marginal R^2^=0.12, p=0.01). Both basal area growth and (log_10_-transformed) branch length were almost completely unrelated to end of season soil moisture (Ψ_PD_), maximum water stress (Ψ_MD_) or ΔΨ, with the exception of branch length marginally significantly increasing with increasing ΔΨ (Fig. 6a,c). Moreover, no water-related climate variables outperformed the null model to explain either growth metric. Indeed, the only climate variables that had lower AICc than the null model were winter/spring temperature-related, and the best climate variable for both radial and stem growth was 30 yr T_min_ (Fig. 6b,d), with sites with warmer winter temperatures growing faster.

**Figure 6:**
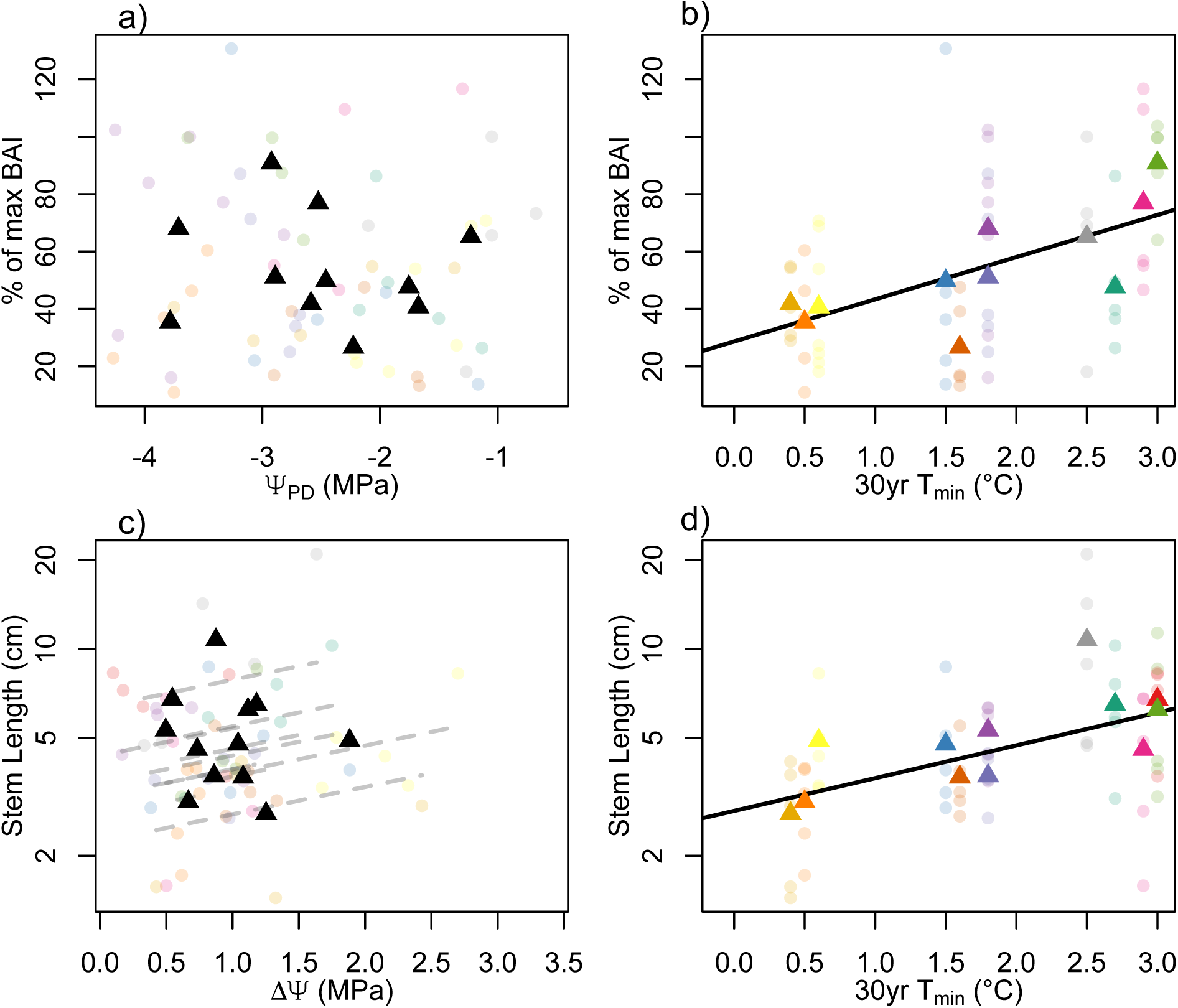
End of season water status does not predict growth, but minimum temperature does. (a) Predawn leaf water potential does not predict radial growth, calculated as the realized % of maximum Basal Area Increment given a tree’s DBH. (b) Radial growth increases strongly with increasing minimum temperatures. (c) Transpirative water potential drop (ΔΨ) is weakly related to average stem length (but Ψ_PD_ and Ψ_MD_ are not, not shown). (d) Stem length is also positively related to site minimum temperature of the coldest month.

Growth was also not related to leaf and stem traits in essentially any expected ways. Most traits were unrelated to either stem or basal area growth, but both within and among sites A_l_:A_s_ and M_l_:M_s_ were both *negatively* related to percent of maximum BAI, and M_l_:M_s_ was also extremely strongly negatively related to branch length (Fig. 6). At the branch level, both A_l_:A_s_ and M_l_:M_s_ decreased strongly with branch length, suggesting a structural, rather than hydraulic or carbon economic driver of both branch allocation traits. If branch length was included as a covariate in models predicting % max BAI, M_l_:M_s_ was no longer significantly related (p=0.21) but A_l_:A_s_ remained significantly negatively related to % max BAI (p=0.01). Across sites, leaf size was positively related to branch length, but not among trees within a site and not to % max BAI. LDMC was marginally significantly related to branch length among trees within sites but not across sites.

### Mechanistic model synthesis

When parameterized with observed, tree-specific A_L_:A_S_ and forced with end-of-season water availability (Ψ_PD_) and meteorology, the HOTTER model simulated Ψ_MD_ that were strongly correlated with but generally more negative than observations (Fig. 8a). Simulated percent loss of conductivity (PLC) was well in line with observed xylem vulnerability (Fig. S5). Variation in VPD did not strongly influence simulated Ψ_MD_ for most trees, but HOTTER predicted that five trees entered a hydraulic death spiral (had very negative Ψ_MD_ and ∼100% loss of hydraulic conductivity) during the median and maximum VPD days of the study period, though not during the lowest VPD day. Simulated end-of-season percent loss of conductivity (PLC) was completely unrelated to observed basal area growth, providing further evidence for a disconnect between end-of-season hydraulic vulnerability and growth (Fig. 8b). Meanwhile, simulated end-of-season gross primary productivity (GPP) was marginally *negatively* correlated with observed growth (Fig. 8c, linear mixed effects model p=0.08). At the same time, net primary productivity, or photosynthesis minus temperature- and allocation-dependent respiration, was negative for all trees but positively related to observed growth (Fig. 8d, p=0.01). Decreasing the leaf area by 50% resulted in no trees entering hydraulic failure and a slightly reduced bias in Ψ_MD_ compared to observations, but remarkably similar results (Fig. S5, S6). Simulations with reduced leaf area also suggested that most trees retained more leaf area (i.e. a higher A_L_:A_S_) than was beneficial based on end-of-season conditions, with both increased (less negative) NPP and decreased PLC in reduced leaf area simulations as well as a reduction in how strongly NPP was negatively correlated with leaf area (Fig. S7).

**Figure 7:**
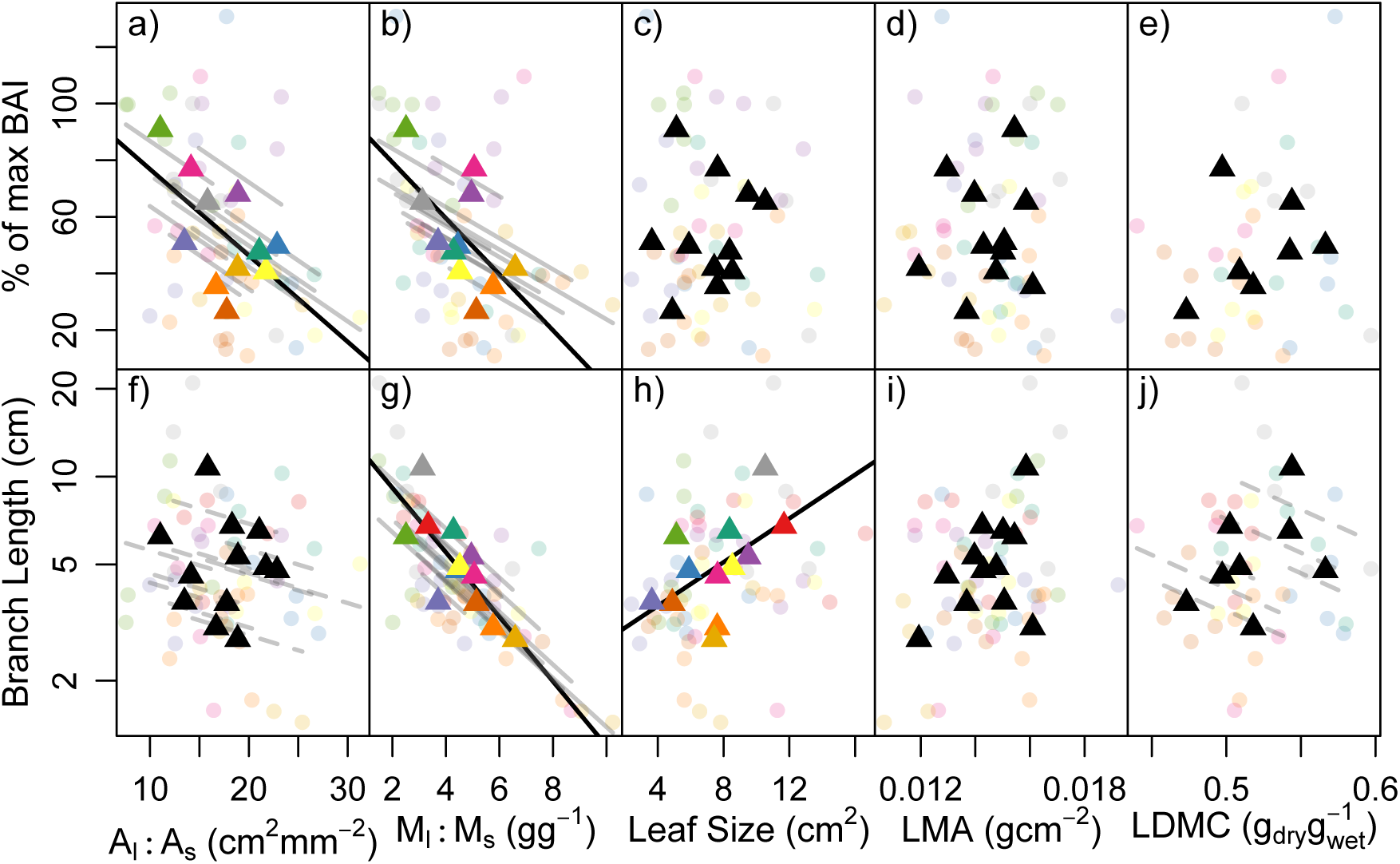
Relationships between leaf traits, including A_l_:A_s_ (a,f), M_l_:M_s_ (b,g), average leaf size (d, h), LMA (d,i), and LDMC (e,j), and radial growth calculated as the percent of realized Basal Area Increment growth (BAI) relative to the 90^th^ percentile growth for that tree size (a-e), and average branch length of the current year terminal branches (1 year of growth, f-j). Points show tree average traits, triangles show site average traits. Gray lines and colored trees show trends among trees within sites, solid/dashed black lines show significant/marginally significant trends from linear mixed effects models (i.e. the mean within-site relationship). Colored site means denote a significant or marginally significant relationship among site means (linear model p<0.1).

**Figure 8:**
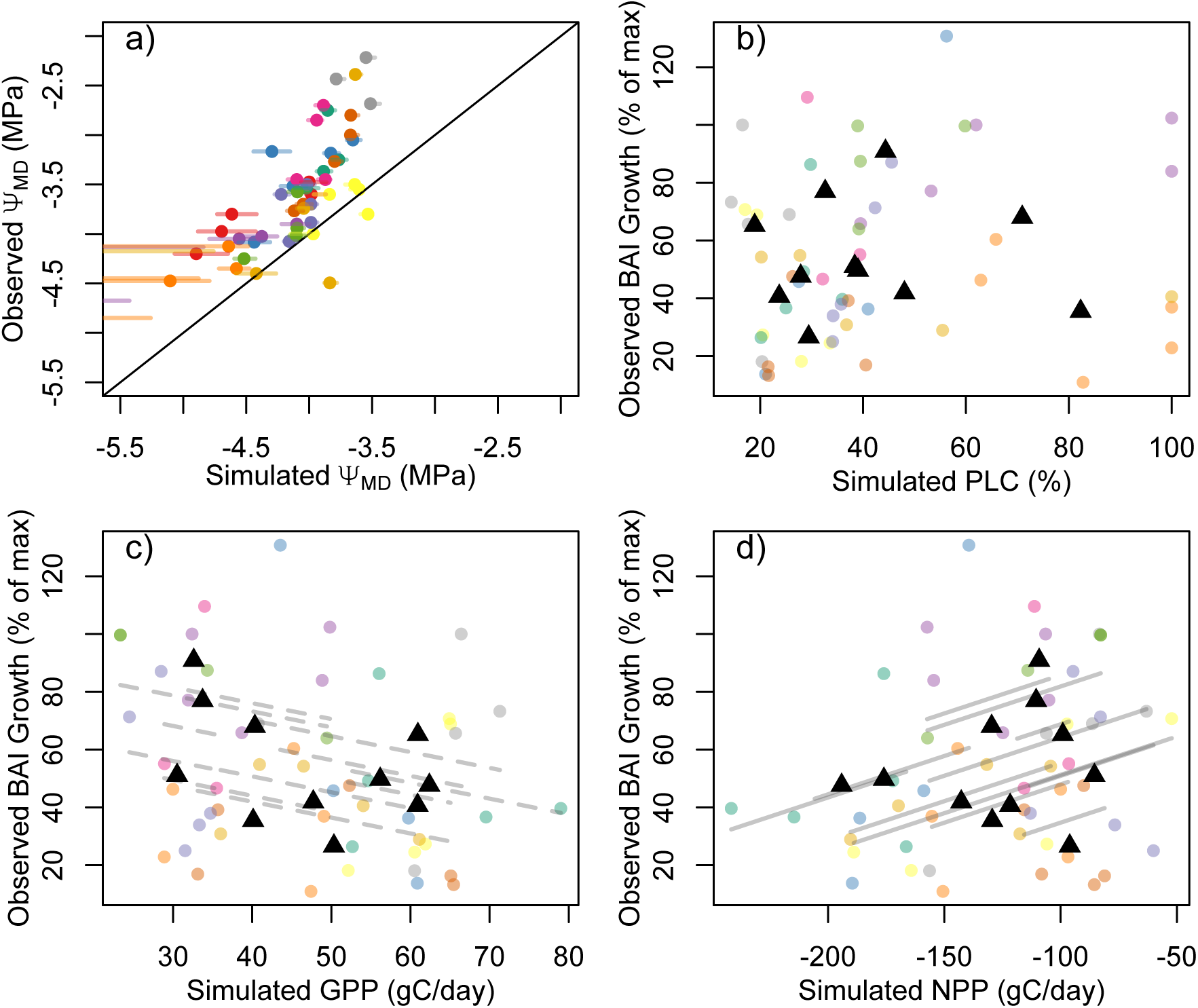
Comparison of variables simulated with the HOTTER mechanistic model and observations demonstrate reasonable model skill and highlight the disconnect between end-of-season hydraulic risk or limitation and observed growth. (a) observed versus simulated midday leaf water potentials (Ψ_MD_), with points showing simulations the median VPD during the sampling (error bars span the lowest VPD day to the highest VPD day during the sampling period). (b) Simulated Percent Loss of Conductivity (PLC) was completely unrelated to observe growth rates (circles = trees, triangles = site means). (c) Simulated gross primary productivity (GPP) was marginally negatively correlated with observed basal area growth. (d) while all trees were simulated to have a negative net primary productivity (i.e. respiration higher than photosynthesis), simulated NPP was significantly positively related to observed growth. (gray lines show trends among trees within sites, dark lines show marginally significant (dashed) or significant (solid) main effects from linear mixed effects models).: Comparison of the optimal A_l_:A_s_ of sampled trees simulated by the HOTTER plant hydraulics model forced with observed soil water potential data (line) to the observed A_l_:A_s_ (points). Trees simulated to have an A_l_:A_s_ (soil water potential <-4 MPa) should be drought deciduous in order to maximize carbon gain (minimize carbon loss).

## Discussion

Our range-wide survey of water availability, water stress, traits and growth in blue oak revealed substantial site-to-site variation in all quantities but remarkable disconnects between them. Site water balance and soil characteristics rarely predicted soil water, water stress, traits, or growth. Moreover, end-of-season water stress, leaf and allocation traits, and growth were largely unrelated to each other, contrary to all our hypotheses. Our only supported hypothesis was that water supply (Ψ_PD_) was more predictable than maximum water stress (Ψ_MD_) due to allocation and stomatal feedbacks. These results raise some difficult questions for both how we predict supply limitation and its effects in deeply rooted tree species in geologically and geomorphically complex landscapes, and how we conceptually integrate hydraulic risk and carbon gain in a highly seasonal environment.

### Deep water access decouples water availability and stress from above-ground climate, making drought stress hard to predict

We found a remarkable decoupling of plant water status from above-ground climate. No single metric of water supply, demand, or balance could explain spatial variation in root-zone soil moisture (Ψ_PD_) or maximum water stress (Ψ_MD_). This finding is in line with historical results showing no relationship between topo-edaphic factors that should govern water availability (slope, aspect, elevation) and blue oak Ψ_PD_ across trees along a ∼3 km elevation gradient (Knops and Koenig 1994) even while year-to-year variation in precipitation drove interannual variation in Ψ_PD_ in the same trees (Knops and Koenig 2000).

The overriding importance of the subsurface, or the ‘weather underground’ (McLaughlin *et al*. 2020), is highlighted by the strong relationship between enriched xylem stable water isotopes and negative plant water potential (both Ψ_PD_ and Ψ_MD_). This partly reflects the influence of meteoric water (Fig. S3b), with precipitation isotopes proving a more successful predictor of Ψ than water balance metrics. In this system, spatial variation in annual precip-weighted δD and δ^18^O is driven by the timing of precipitation and storm size, with more isotopically depleted sites receiving a larger fraction of rain as large, cold, winter storms. Coincidentally, these same types of storms that are likely to infiltrate to recharge subsurface water pools such as shallow aquifers and fractured bedrock, providing deep, isotopically depleted water for blue oaks to use far into the growing season. Moisture balance metrics evidently did not capture this hydrological dynamic.

However, even when controlling for the meteoric water background, more enriched sites and more enriched trees at a site tended to have more negative Ψ_PD_. The large geographic scope of this project precluded the characterization of isotopic end members for each site, but isotopic enrichment likely indicates increased reliance on evaporatively enriched soil moisture from the shallow soil, while relatively isotopically depleted xylem water reflects access to deeper moisture sources (ground water, rock water) (Ehleringer *et al*. 1993; Hahm *et al*. 2020; Sprenger *et al*. 2016). The absolute functional rooting depth implied by these isotopic values may vary based on site hydrology, particularly if regolith properties (e.g. highly rocky substrates) govern the depth of soil evaporative enrichment. However, the main driver of variation in end-of-season water supply, and consequently maximum water stress, appears to be access to relatively deep water sources.

### Growth and hydraulic risk are decoupled in a Mediterranean-type climate

Water stress clearly limits assimilation in blue oak across much of its range by the end of the growing season. Leaf-level gas exchange showed a strong threshold decline as leaf water potential approached the point of substantial leaf embolism (leaf P50 measured using the optical technique (Skelton *et al*. 2019)). This decline was consistent in midday gas exchange rates across sites, within a single tree over the course of a morning (Fig. 4), and previously at a sight over the course of a growing season (Xu & Baldocchi 2003). Such behavior is highly consistent with our understanding of hydraulic cost-driven stomatal behavior (Anderegg *et al*. 2018; Sperry *et al*. 2016). Indeed, many sampled trees appeared quite close to the hydraulic edge, with a number of trees surpassing leaf and even stem P50 at midday (Fig. 2) and five trees simulated to have nearly 100% loss of hydraulic conductivity without altering VPD or leaf area in the HOTTER model (Fig. 8, Fig. S5,S6,S7).

If end-of-season water availability is correlated with conditions earlier in the growing season, we might assume that tree growth would decline across sites as whole plant carbon gain becomes increasingly constrained by negative leaf water potentials, causing a negative relationship between growth and late season hydraulic damage. This was not the case (Fig. 6). Indeed, simulated end-of-season hydraulic damage and gross photosynthesis from a coupled photosynthesis and plant hydraulics model had no relationship (PLC, Fig. 8b) or a negative relationship (GPP, Fig. 6c) with observed radial growth.

One explanation for this disconnect is a temporal mismatch between the times of year most critical for carbon gain/growth and those that determine hydraulic risk from drought stress. Due to California’s Mediterranean-type climate (cool wet winters and hot dry summers), photosynthesis in blue oak woodlands peaks in the spring and early summer, up to 50 days earlier than solar radiation peaks (Ma *et al*. 2011; Xu & Baldocchi 2003). Photosynthetic capacity and dark respiration both decline precipitously less than 40 days after leaf out in blue oak, with dark respiration in particular becoming quite low through the majority of the mid and late summer (Xu & Baldocchi 2003). Thus, the most productive growing season for blue oaks is a small fraction of their total leaf-on period. Our results suggest that the water status dynamics of the later growing season, which determine blue oak’s proximity to hydraulic damage thresholds, may be fundamentally disconnected from the early growing season factors that drive carbon gain and growth.

A second temporal disconnect that may decouple growth from end-of-season water stress is the use of prior year(s) carbon to fuel current year growth. Lagged growth effects, memory effects or legacy effects are ubiquitous in tree rings (Anderegg *et al*. 2015; Klesse *et al*. 2023; Ogle *et al*. 2014). The use of prior year carbon is hypothesized to underpin a disconnect between flux tower estimates of annual gross primary productivity and tree ring estimates of net primary productivity (Cabon et al. 2022). In blue oak at dry sites and during dry years, turgor-driven limits to radial growth likely manifest early in the growing season (when predawn water potentials drop below -0.8 MPa for many plants, (Körner 2015; Potkay *et al*. 2022)). Consequently, the carbon fixed over the bulk of the leaf-on period will support respiration and future growth, rather than current year growth. Thus, the end-of-season water limitation we document most likely influences future growth.

Finally, our HOTTER simulations suggest a larger role for respiration than hydraulic limitations to assimilation in determining late growing season carbon balance. All trees had a simulated negative net carbon balance during the sampling period (i.e. were respiring more than they photosynthesized), but simulated NPP was still positively correlated with observed growth, even while simulated GPP was negatively correlated with growth (Fig. 8). This could suggest that, even under extreme water limitation, temperature- and allocation-driven respiration demand may still be more important for spatial variation in blue oak growth than hydraulic damage.

Our results also raise a subsidiary question. Why do the spatial controls on blue oak growth not mirror the inter-annual controls on growth? Blue oak is a canonical dendroclimatology and dendro-hydrology species (Stahle 2013), often considered ‘better than rain gauges’ for tracking interannual variation in precipitation. Yet spatial variation in mean growth was not related to either end-of-season water availability or water-related climate/meteorological variables. Instead, spatial variation in growth was correlated with minimum annual temperature of the coldest month (T_min_), suggesting that other plant processes related to winter temperature, such as the onset of spring leaf out, may control site-to-site variation in growth. A contrast between temporal and spatial growth sensitivity to climate has recently been documented in multiple North American tree species (Canham *et al*. 2018; Klesse *et al*. 2020; Perret *et al*. 2024). This implies that climatically-driven interannual variation in tree growth is disconnected from the climatic drivers of spatial variation in mean growth among populations because of large local adaptation or acclimation (Canham *et al*. 2018). Interannually, no adaptive response is possible and many traits (rooting depth, whole tree acclimation) are slow to respond plastically, making the phenotype relatively fixed. Meanwhile, extensive phenotypic adjustments, both plastic and ecotypic, can mediate the growth impacts of spatial variation in climate. This appears to strongly be the case for blue oaks, and is supported by the large among-site variation in leaf and allocation traits (Table S1), hydraulic traits other than P50 (Anderegg *et al*. 2023) and ecotypic variation in seedling performance (McLaughlin *et al*. 2022).

### Traits have complicated cause-vs-effect relationships with growth and hydraulic risk

Leaf traits were generally unrelated to end of season plant water potentials (Fig. S4), water balance-related climate variables (Fig. 5), and growth rates (Fig. 7). LDMC was the only trait related to end-of-season water potential, surprisingly increasing with less negative Ψ_MD_ (Fig. S4, mixed effects model p=0.005) in a pattern that warrants further study. Meanwhile, leaf size was the only trait correlated with site climate (LMA and A_L_:A_S_ were related to soil water holding capacity but not significantly related to any climate predictor). The decrease in leaf size at drier sites across the landscape mirrors the among-population genetic differentiation in leaf size previously seen in a blue oak common garden (Anderegg *et al*. 2023), and may represent an adaptive response to drought stress. Decreased leaf size can decrease the maximum leaf temperatures experienced in hot parts of blue oak’s range when transpiration is limited (Baldocchi & Xu 2007). This trend is also mirrored across oak species, with drier-adapted oaks generally having smaller leaves (Skelton et al. 2021).

However, beyond the putatively adaptive variation of leaf size, few trait-climate, trait-water stress, or trait-growth relationships were clear or consistent. Leaf to stem allocation, either expressed as the hydraulic demand relative to transport capacity A_l_:A_s_ or as the carbon economic M_l_:M_s_, was strongly *negatively* related to growth rates. This directly contrasts with theory (Lambers & Poorter 1992) and observations in herbs (Poorter et al. 2012), which find that maximizing investment in leaves should pay compounding interest and maximize Relative Growth Rates. Instead, it suggests that leaf to stem allocation, at the branch level, may be the *consequence* rather than the cause of growth rate variation in blue oak. Trees with high net carbon uptake can afford to grow more structural tissue, and thus produce longer branches with more stem mass per leaf area. For A_l_:A_s_, branch-level trait values are jointly driven by the positive influence of leaf size (larger leaves drive higher A_l_:A_s_) and the negative influence of branch length (longer branches produce lower A_l_:A_s_), and leaf size and branch length are uncorrelated (Fig. S8). Thus, some of the counter-intuitive A_l_:A_s_ relationship with basal area growth is an artifact of branch length. However, even controlling for branch length (either by including it as a covariate or analyzing the raw residuals of the log(A_l_:A_s_) ∼ log(branch length) relationship), A_l_:A_s_ retains a significant negative relationship with BAI (p=0.009 for both approaches). Collectively, this could support A_L_:A_S_ being a *consequence* of a tree’s carbon balance from the prior years.

However, these results may also suggest that A_L_:A_S_ *causes* growth variation, just in the wrong direction due to structural overshoot. Leaf size and A_L_:A_S_ were positively correlated with growth rates in a mesic common garden experiment (Anderegg et al. 2023), but negatively correlated (A_l_:A_s_) or uncorrelated (leaf size) with growth in the wild. The 2000-2021 period was the driest 22 year period in at least the last 1200 years in the Southwestern U.S. (Williams et al. 2022), and the sampling year was much drier than average (Fig. 1). Against this megadrought background, large leaf area or A_L_:A_S_ may have actually been maladaptive over the study period driving reduced growth in many sites, even if it generally promotes growth in a mesic common garden.

Ultimately, the large site-to-site variation in all traits suggests that traits are responding to a complex combination of environmental cues and endogenous factors (e.g. growth rate) that we still don’t fully understand. In functional ecology, we often focus on the consequences of traits for fitness components such as growth or survival (Violle *et al*. 2007). However, our results highlight that, particularly within species, trait variation can be challenging to interpret, as it represents (presumably) adaptive responses to maximize sometimes orthogonal fitness components. For instance, in blue oak, leaf morphology and allocation to leaves versus stems may be simultaneously trying to optimize early growing season carbon gain and growth and minimize end of growing season mortality risk due to hydraulic damage, which we have found to be largely unrelated. Add in the feedback effects of growth rate on traits themselves and the difficulty of disentangling cause versus effect becomes extremely challenging. The mechanistic modeling application presented here hopefully lays the foundation for future exploration of this complex challenge.

## Conclusion

We conducted a range-wide survey of blue oak supply limitation and drought stress at the end of a substantial drought year, visiting 15 sites over an area 650 km north to south. We found a remarkable disconnect between site water balance, end-of-season drought stress, growth, and drought avoidance-related allocation traits. These results highlight our profound uncertainty about the edaphic environment of deeply rooted tree species - stem water isotope composition strongly predicted leaf water potentials but multiple soils data sources did not. Our results also highlight how, particularly in Mediterranean-type climates, the temporal mismatch between water stress and the early season conditions that determine growth and allocation traits challenges our simple assumptions about maximizing carbon gain and organismal performance under water limitation. Drought stress and its consequences are fundamentally hard to predict across the landscape, strongly challenging our ability to predict the spatial patterns of future drought-induced forest mortality.

## Acknowledgements

We acknowledge the Traditional Custodians and Owners of California, and recognize their continuing connection to land upon which this research was conducted. We particularly recognize the People’s on whose traditional territory our labs and study sites sit, including the Chumash, Sierra Miwok, Konkow, Pomo, and Northern Wintu. We acknowledge their Elders both past and present, and their future generations. We thank the Hopland Research Extension, the Ranger Station at Sonora, and the San Juaquin Experimental Range for access to trees on their property. We also acknowledge funding from the National Science Foundations (NSF 1457400 to DDA & TED; NSF DBI-1711243 to LDLA; 2003205, and 2216855 to LDLA and ATT), the National Oceanographic and Atmospheric Administration (Climate and Global Change Fellowship to LDLA), and the California Board of Forestry and Fire Protection (CALFIRE, grant 8GG21813 and 8GG23808 to LDLA). ATT acknowledges funding from University of California Laboratory Fees Research Program Award No. LFR-20-652467, and the Gordon and Betty Moore Foundation GBMF11974.

## Author Contributions

LDLA, RPS, PP, DDA and TED conceived of the project ideas; LDLA, RPS, PP and ATT designed the methodology; LDLA, RPS, JD and PL collected the data; ATT performed model simulations; LDLA analyzed the data and lead the writing of the manuscript. All authors contributed to editing the manuscript.

