## Supplementary material for "Maximum water stress is decoupled from climate, traits and growth in a xeric oak": Fig. S1

1 – UC Santa Barbara, Ecology, Evolution & Marine Biology

Address: Ecology, Evolution & Marine Biology
University of California, Santa Barbara
Santa Barbara, CA 93106-9620

541.790.1096

2 – UC Berkeley, Integrative Biology, Berkeley, CA, USA

3 – South African Environmental Observation Network and School of Animal, Plant and Environmental Sciences, University of Witwatersrand, Johannesburg, South Africa

4 – UC Santa Barbara, Geography, Santa Barbara, CA, USA

5 – UC Berkeley, Environmental Science, Policy & Management, Berkeley, CA, USA

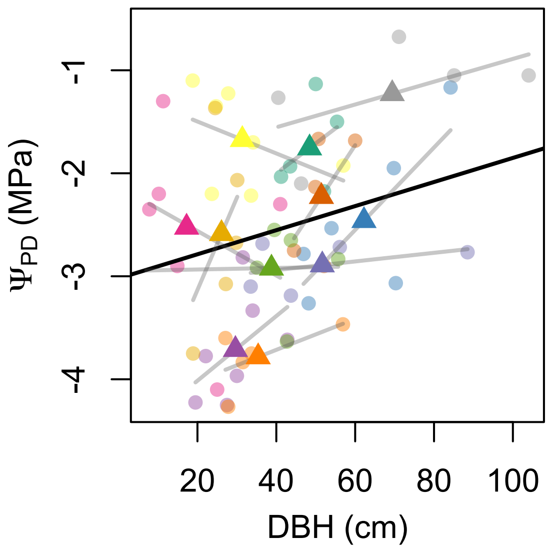


**Figure S1:** *Predawn water potential ($\Psi_{PD}$) was generally less negative in trees with larger diameter at breast height (DBH), suggesting that larger trees had access to increased soil moisture. Points show tree average values, triangles show site average values, colors indicate different sites, and gray lines show the trend among trees at a site. Black trend line show statistically significant ($p<0.05$) fixed effects from a mixed model with site random intercepts.*

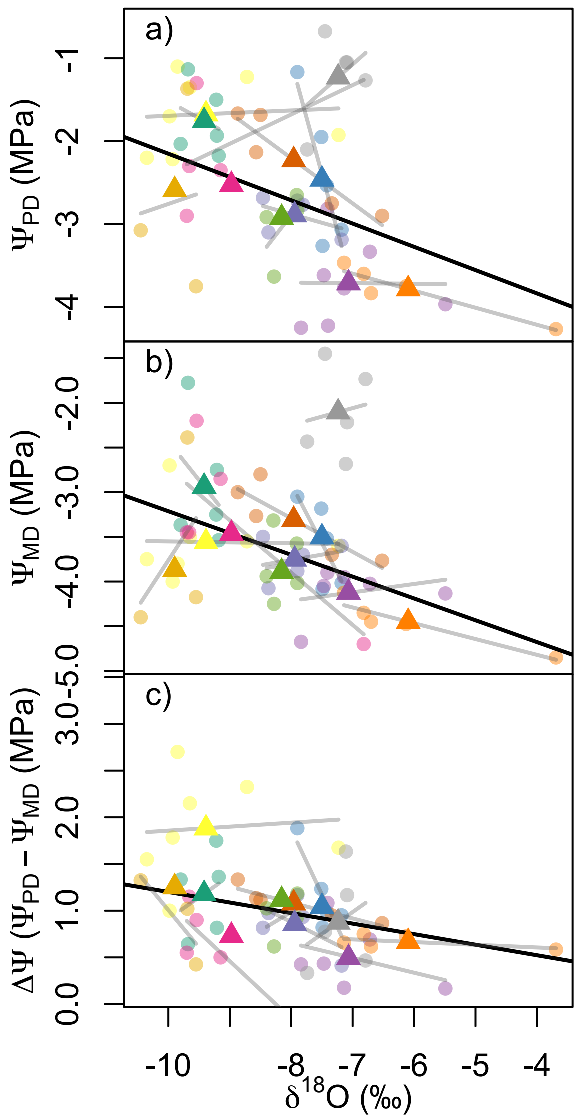


**Figure S2:** *Plant water potentials were strongly related to xylem water isotopes within and among sites (similar to Fig. 3 but with $\delta^{18}O$). Predawn leaf water potential (a), midday leaf water potential (b) and the difference between them (c) decrease with increasing xylem water enrichment of $\delta^{18}O$. Points show tree average values, triangles show site average values, colors indicate different sites, and gray lines show the trend among trees at a site. Black trend line show statistically significant ($p<0.05$) fixed effects from a mixed model with site random intercepts, with the outlier burned site (gray points, triangle) excluded.*

**Table S1:** Summary of among-site variability in water status, traits and growth.

| TRAIT | N | ETA2 | OMEGA2 | AMONG SITE P-VALUE | CV | BEST ENVIRONMENTAL PREDICTOR | ENV PREDICTOR P | ENV VARIANCE EXPLAINED |
|---|---|---|---|---|---|---|---|---|
| **PD** | 83 | 0.76 | 0.71 | 0 | 0.46 | 30yr AET | 0.108 | 0.132 |
| **MD** | 86 | 0.6 | 0.52 | 0 | 0.23 | WY PPT 2018 | 0.191 | 0.074 |
| **E.DROP** | 82 | 0.66 | 0.58 | 0 | 0.59 | GS VPD 2018 | 0.011 | 0.245 |
| **AL_AS** | 60 | 0.45 | 0.32 | 0.001 | 0.29 | 1m Soil H2O Storage | 0.02 | 0.187 |
| **ML_MS** | 60 | 0.33 | 0.17 | 0.037 | 0.44 | Tmin 2018 | 0.012 | 0.176 |
| **LEAFSIZE** | 60 | 0.51 | 0.39 | 0 | 0.43 | 30yr PPT | 0.026 | 0.197 |
| **LMA** | 60 | 0.48 | 0.35 | 0 | 0.12 | PAW | 0.019 | 0.201 |
| **LDMC** | 40 | 0.66 | 0.58 | 0 | 0.07 | SP Tmax 2018 | 0.116 | 0.212 |
| **LENGTH** | 60 | 0.43 | 0.3 | 0.002 | 0.35 | Tmin 2018 | 0.043 | 0.145 |
| **PERC_MAXBAI** | 59 | 0.35 | 0.21 | 0.014 | 0.57 | 30yr Tmin | 0.01 | 0.186 |

**Table S2**: Model results predicting leaf water potential from xylem water stable isotopes. p: p-value of isotope predictor in linear mixed effect model with site as random effect. R2m: marginal $R^2$ (considering fixed effects); R2c: conditional $R^2$ (including both fixed and random effects) from linear mixed effects models. 'btw site R2': the amount of among-site variation in water potential explained by isotopic predictor based on linear model of site-averaged data.

| | $\delta D$ | | | | $\delta^{18}O$ | | | |
|---|---|---|---|---|---|---|---|---|
| All data | p | R2m | R2c | btw site R2 | p | R2m | R2c | btw site R2 |
| $\Psi_{PD}$ | **0.012** | **0.181** | **0.672** | **0.216** | **0.046** | **0.08** | **0.634** | **0.223** |
| $\Psi_{MD}$ | **0.015** | **0.154** | **0.669** | **0.012** | **0.028** | **0.093** | **0.581** | **0.033** |
| E.drop | **0.006** | **0.23** | **0.389** | **0.575** | 0.065 | 0.086 | 0.42 | 0.419 |
| no PWD outlier | | | | | | | | |
| $\Psi_{PD}$ | **0** | **0.605** | **0.636** | **0.829** | **0.004** | **0.21** | **0.536** | **0.573** |
| $\Psi_{MD}$ | **0.001** | **0.362** | **0.469** | **0.498** | **0.001** | **0.271** | **0.346** | **0.356** |
| E.drop | **0.003** | **0.297** | **0.455** | **0.612** | 0.06 | 0.094 | 0.478 | 0.409 |

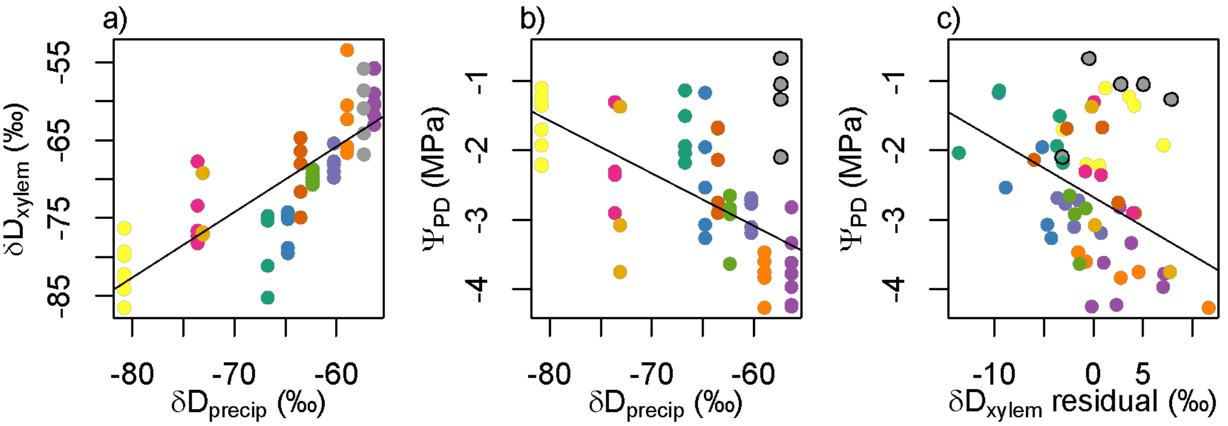


**Figure S3:** The isotopic values of meteoric water ($\delta D_{precip}$) are positively related to the $\delta D$ of tree xylem water (a), and negatively related to soil water availability ($\Psi_{PD}$, b). However, the residuals of the $\delta D_{xylem}$~$\delta D_{precip}$ relationship (i.e. how enriched xylem is relative to a site's precipitation-weighted average meteoric context) still predicted soil moisture availability (c), with more relatively enriched xylem water being associated with more negative $\Psi_{PD}$. Patterns with $\delta^{18}O$ are qualitatively similar to those shown with $\delta D$.

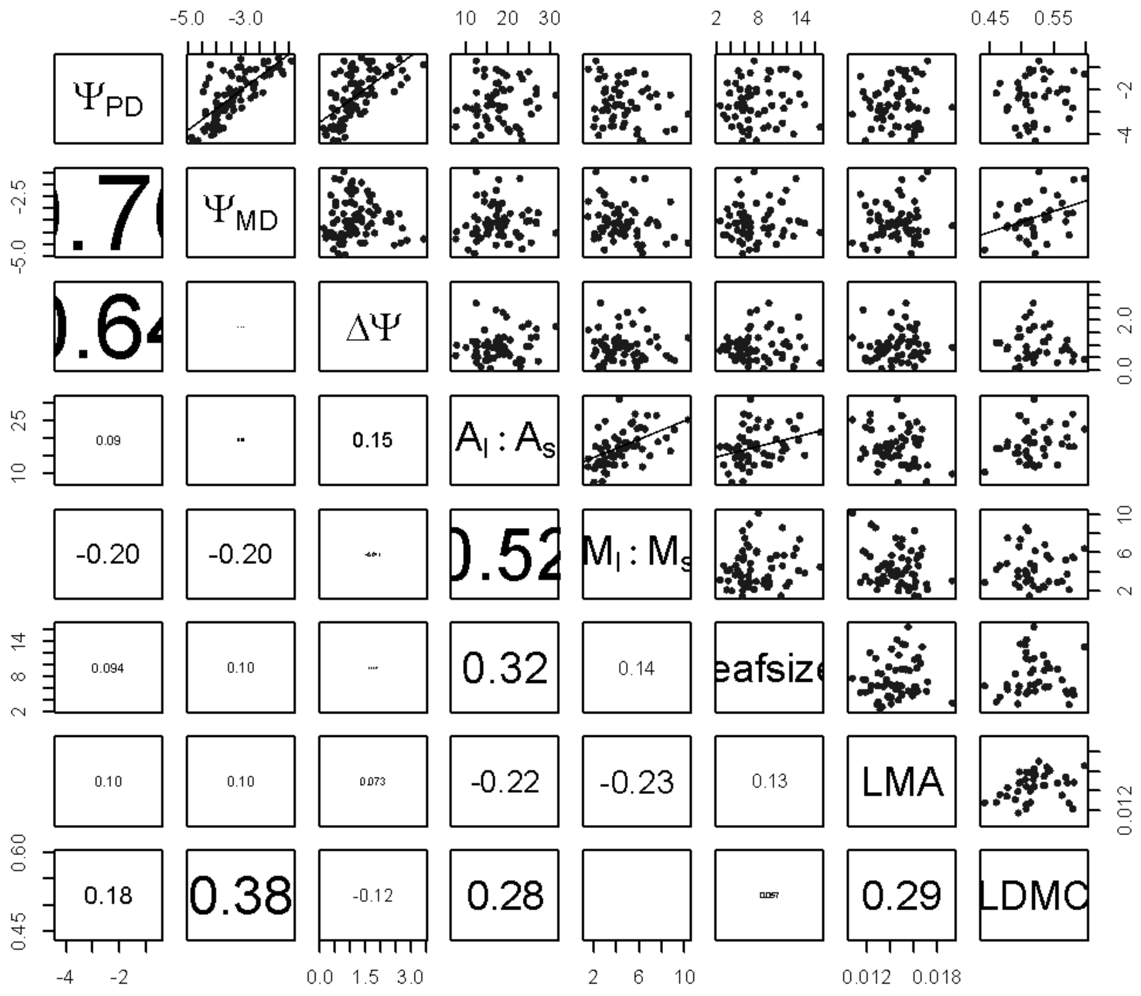


**Figure S4:** Pair plots of water potentials and leaf and allocation traits. Upper diagonal shows scatterplots between variables (significant correlations p<0.5 with trend lines) and lower diagonal shows correlation coefficients (text scaled by correlation absolute value).

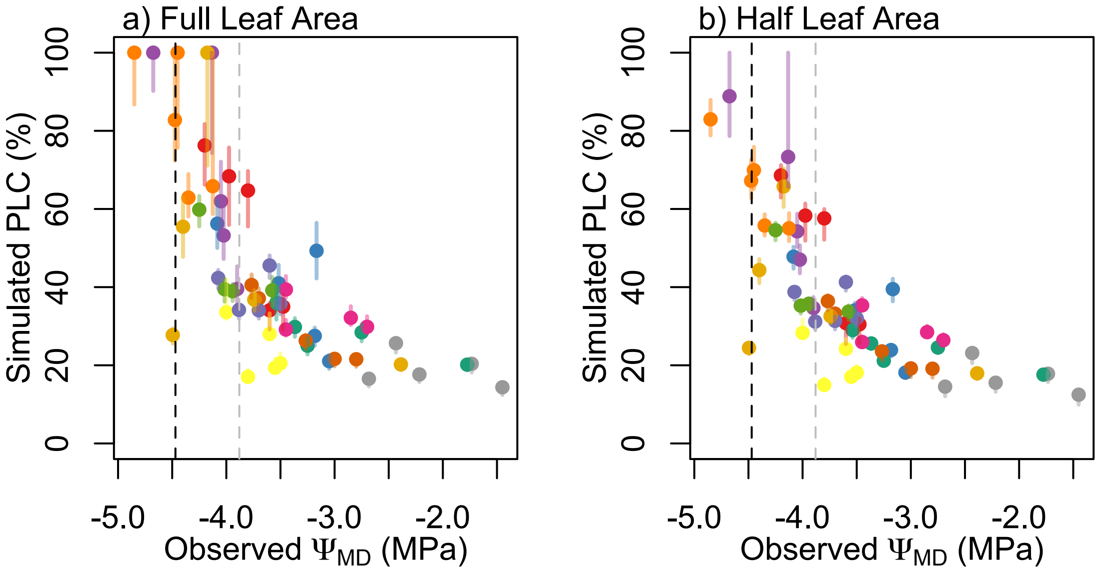


**Figure S5:** (a) Simulated whole-plant percent loss of conductivity (PLC) as a function of observed $\Psi_{MD}$, with leaf (gray) and stem (black) P50 as dotted lines. (b) Identical simulations but with tree leaf area reduced by 50%. Point colors indicate sites, error bars indicate variation from the highest VPD to lowest VPD day during the sampling period, points show mean VPD simulations.

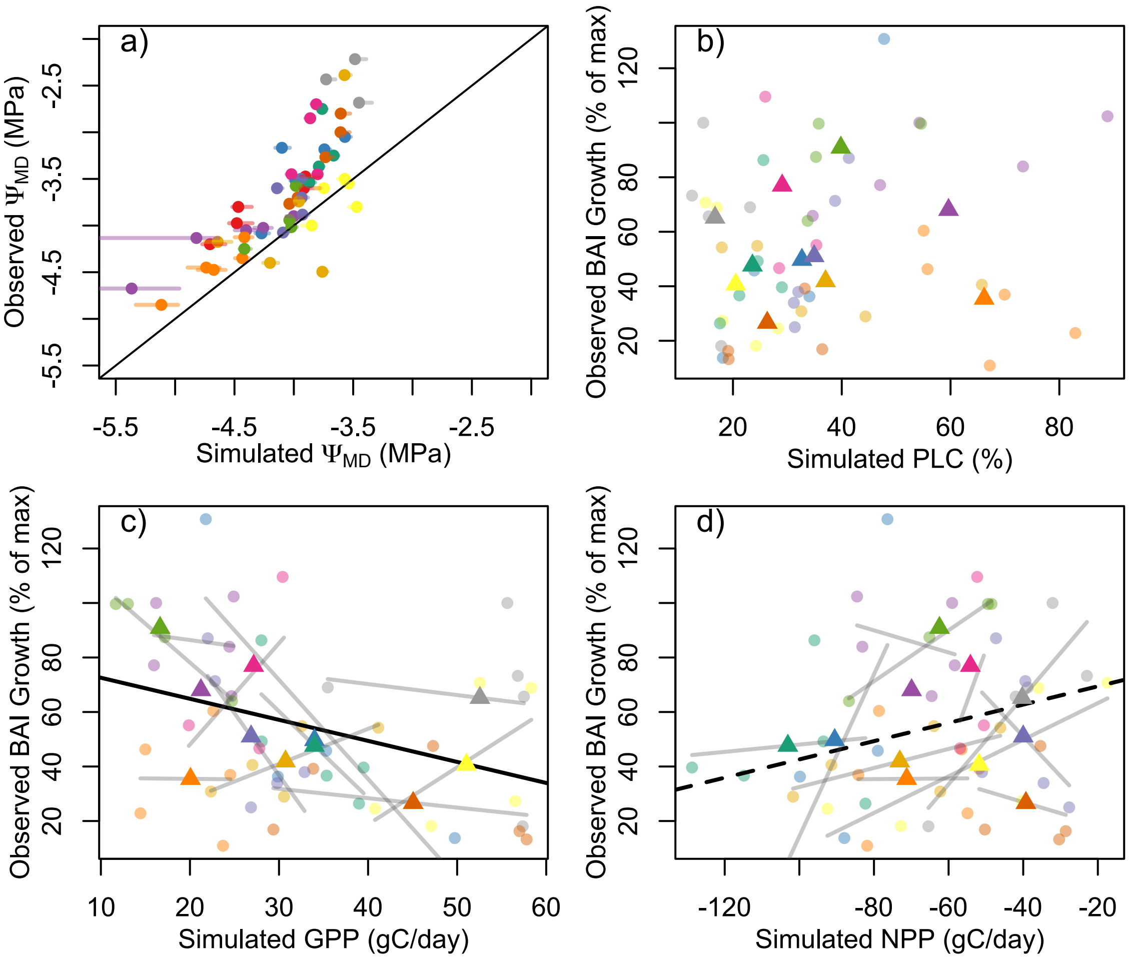


**Figure S6:** Comparison of variables simulated with the HOTTER mechanistic model, decreasing whole-tree leaf area by 50% compared to Fig. 8, and observations. (a) observed versus simulated midday leaf water potentials ($\Psi_{MD}$), with points showing simulations the median VPD during the sampling (error bars span the lowest VPD day to the highest VPD day during the sampling period). (b) Simulated Percent Loss of Conductivity (PLC) was completely unrelated to observe growth rates (circles = trees, triangles = site means). (c) Simulated gross primary productivity (GPP) was marginally negatively correlated with observed basal area growth. (d) while all trees were simulated to have a negative net primary productivity (i.e. respiration higher than photosynthesis), simulated NPP was significantly positively related to observed growth. (gray lines show trends among trees within sites, dark lines show marginally significant (dashed) or significant (solid) main effects from linear mixed effects models).

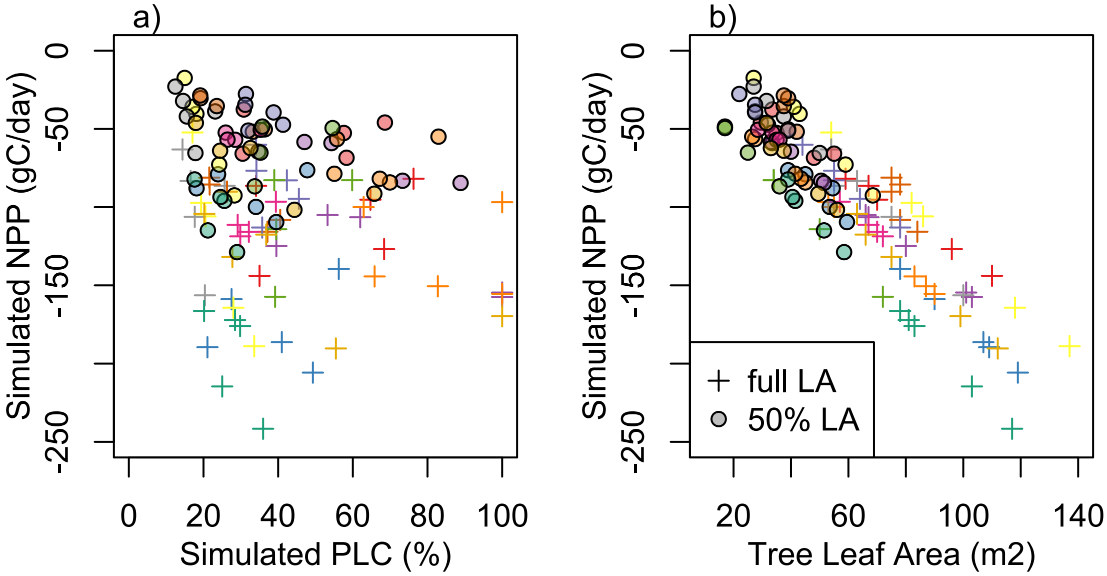


**Figure S7:** (a) HOTTER simulated whole-plant percent loss of conductivity (PLC) and net primary productivity (NPP), illustrating the difference between simulations with full tree leaf area (crosses) and simulations decreasing leaf area by 50% (points) (b) The extent of negative NPP (carbon loss through respiration) is proportional to the allocation to leaf area control simulations, but becomes less sensitive to leaf area in 50% leaf area simulations when more trees retain open stomata.

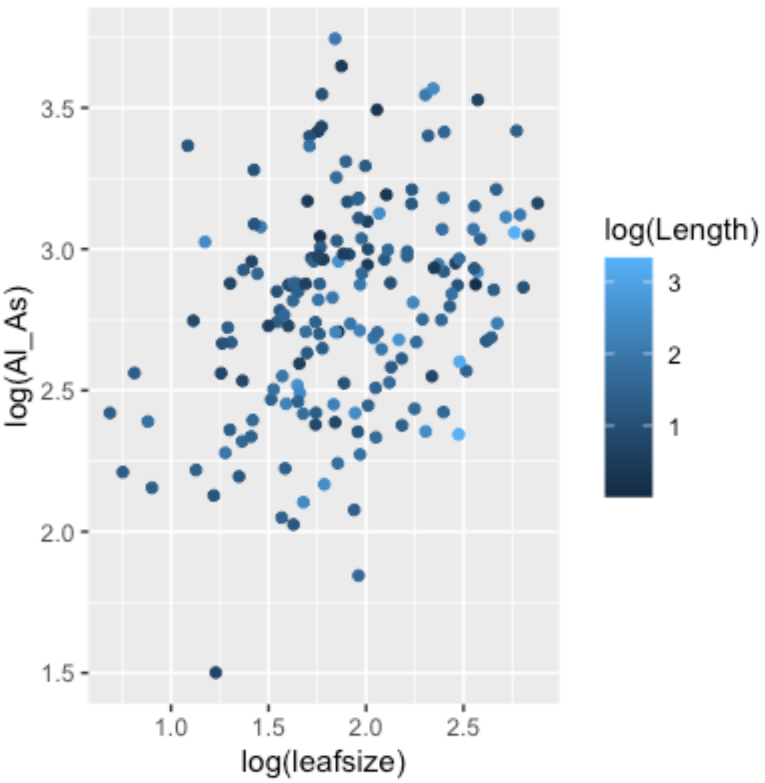


**Figure S8:** $A_l$:$A_s$ at the branch level is jointly controlled by leaf size (larger leaves produce higher $A_l$:$A_s$, linear mixed effects model with site and tree random intercept $p<0.0001$) and branch length (longer branches produce lower $A_l$:$A_s$, $p=0.006$).

**Table S3:** Model output from linear mixed effects model relating log($A_L$:$A_S$) to log(leaf size) and log(branch length), with a random intercept for tree and site. There was no significant interaction between branch length and leaf size, based on a Likelihood Ratio Test.

| | Estimate | Std. Error | df | T value | P value |
|---|---|---|---|---|---|
| Intercept | 2.28783 | 0.15343 | 91.55937 | 14.911 | < 2e-16 *** |
| log(length) | -0.12612 | 0.04503 | 162.95509 | -2.801 | 0.00572 ** |
| log(leafsize) | 0.36052 | 0.07036 | 122.67828 | 5.124 | 1.13e-06 *** |
